## Supplementary Figures and Table for "Chemical engineering of therapeutic siRNAs for allele-specific gene silencing *in vivo* in CNS"

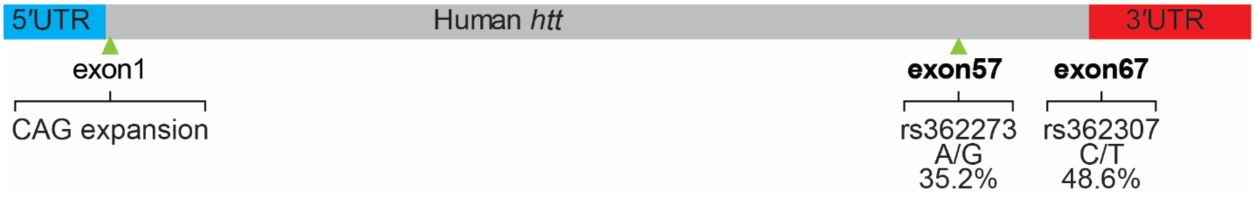


***Supplementary Figure 1.*** A simplified map of the human *htt* gene shows regions of interest. Exons which are highlighted in bold contain SNPs with high rates of heterozygosity, used for subsequent experiments. 35.2% of HD patients are heterozygous (A/G) at rs362273, and 48.6% of HD patients are heterozygous (C/T) at rs362307[1,2].

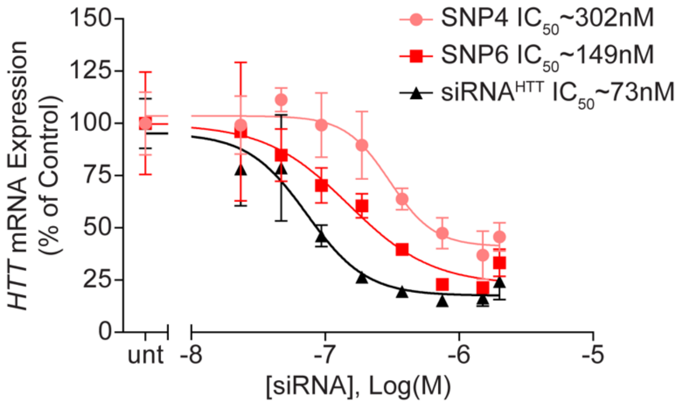


***Supplementary Figure 2.*** **SNP-targeting siRNAs silence htt mRNA with a moderate reduction in potency compared to pan-HTT targeting siRNAs.** HeLa cells were treated with siRNAs via passive uptake for 72 hours. Endogenous huntingtin mRNA levels were measured with Quantigene 2.0 assay, and normalized to *PPIB*.

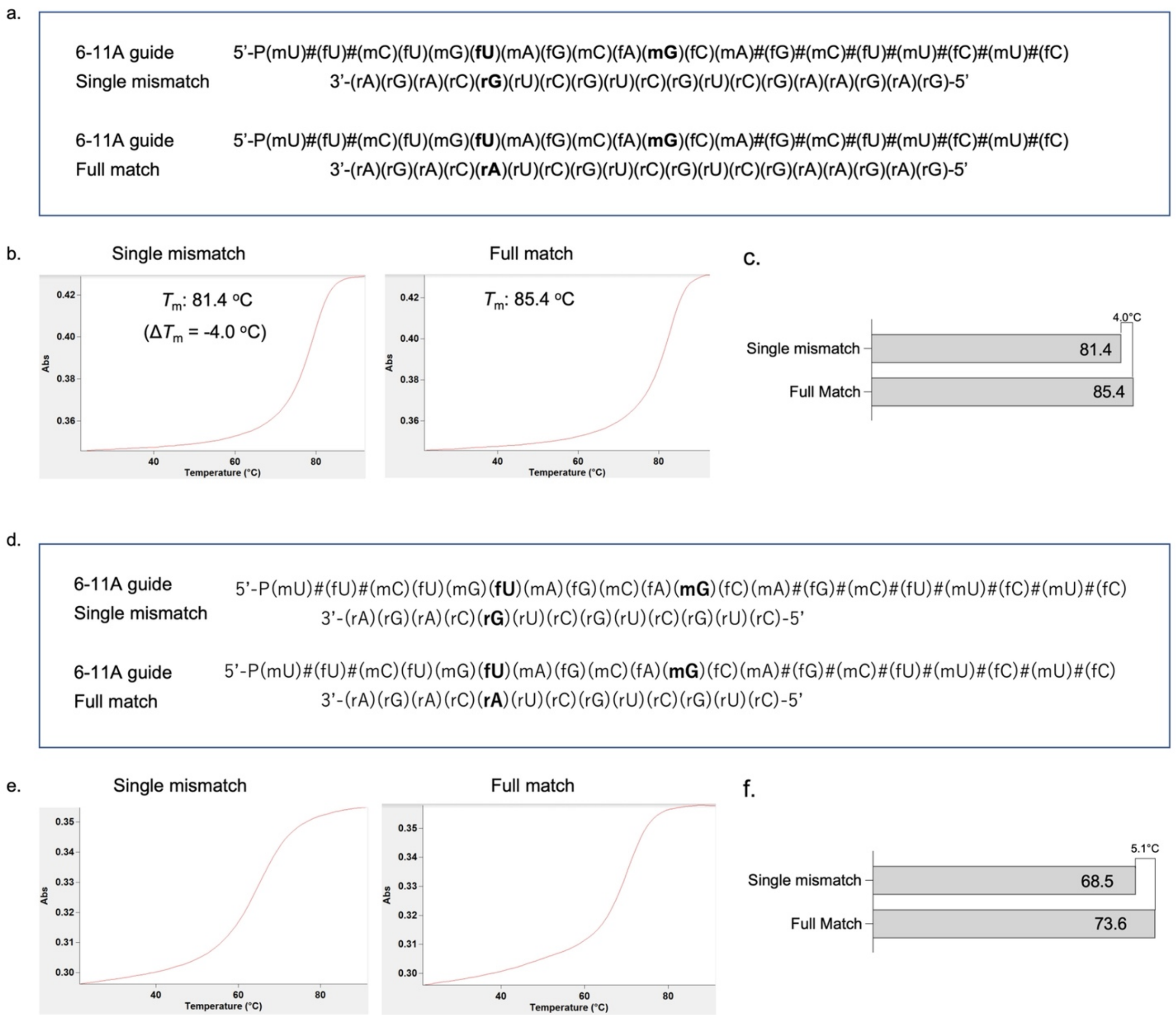


***Supplementary Figure 3*****: A single mismatch in the seed region of the siRNA guide strand does not have a significant impact on duplex stability.** Guide strands for SNP6-11 were hybridized to 19 nucleotide and 13 nucleotide RNA strands, with or without a mismatch in the seed, and melting temperature was measured by thermo stability assay. **(a,d)** Sequence and structureofthe 6-11 guide strand and a complementary 19 nucleotide RNA or 13 nucleotide RNA. **(b,e)** Tm curves for SNP6-11 hybridized to a 19mer complementary RNA or 13mer complementary RNA strand, with a single mismatch or full matched sequence. **(c,f**) Graph comparing melting temperatures of the full match sequence with a single mismatch included, exhibiting a minimal change in melting temperature, whether hybridized to a 13nt or 19nt complementary RNA strand.

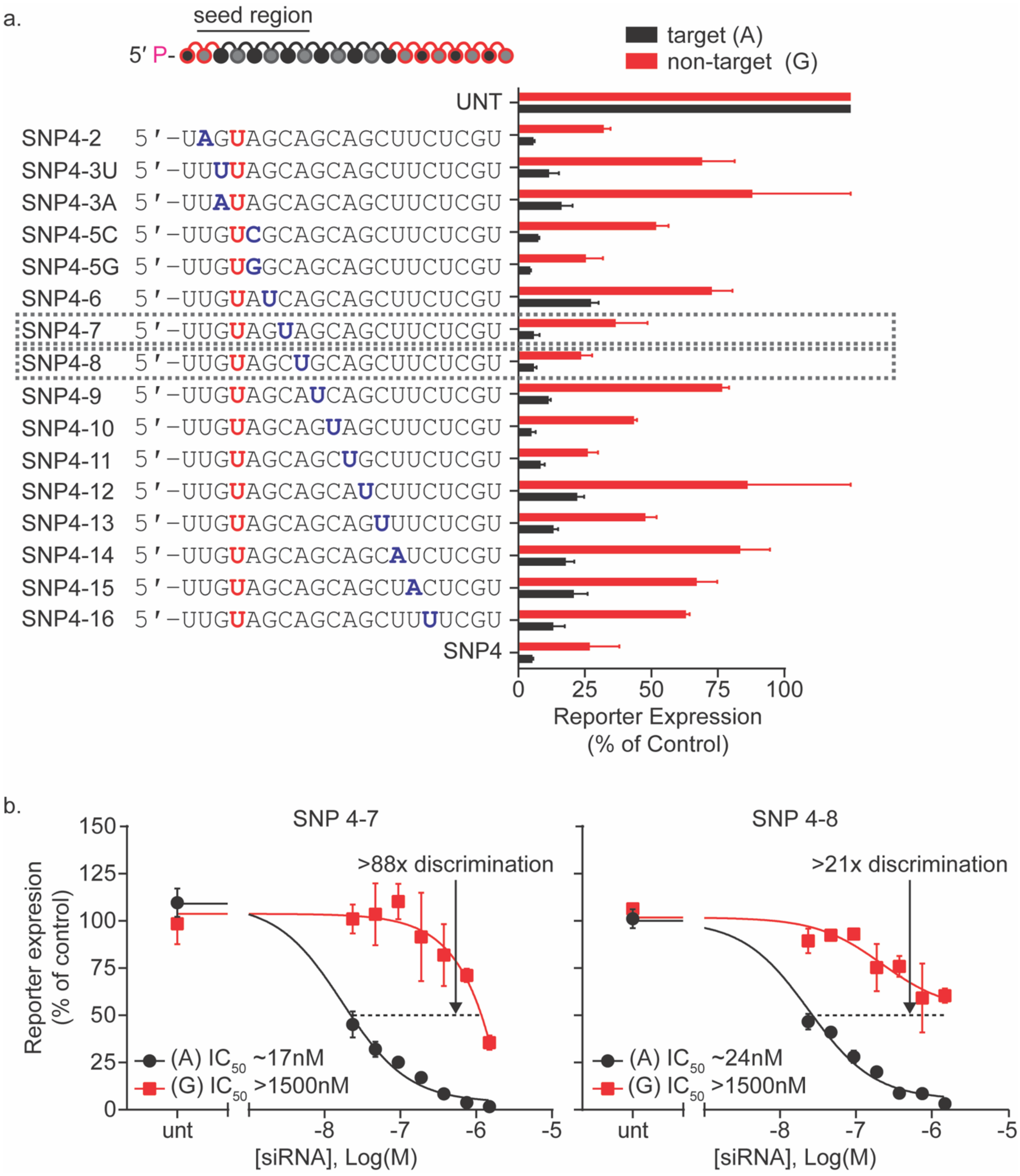


***Supplementary Figure 4*. Addition of secondary mismatch is well tolerated in target silencing, but reduces non-target activity. (a)** siRNA SNP4 (from primary screen; main Fig. 1) was screened for introduction of secondary mismatch, resulting in increased discrimination. **(b)** Dual-luciferase reporter assay dose response of lead compounds identifies SNP4-7 as the best-performing siRNA, improving discrimination more than 10x when compared to SNP4 (main Figure 1c).

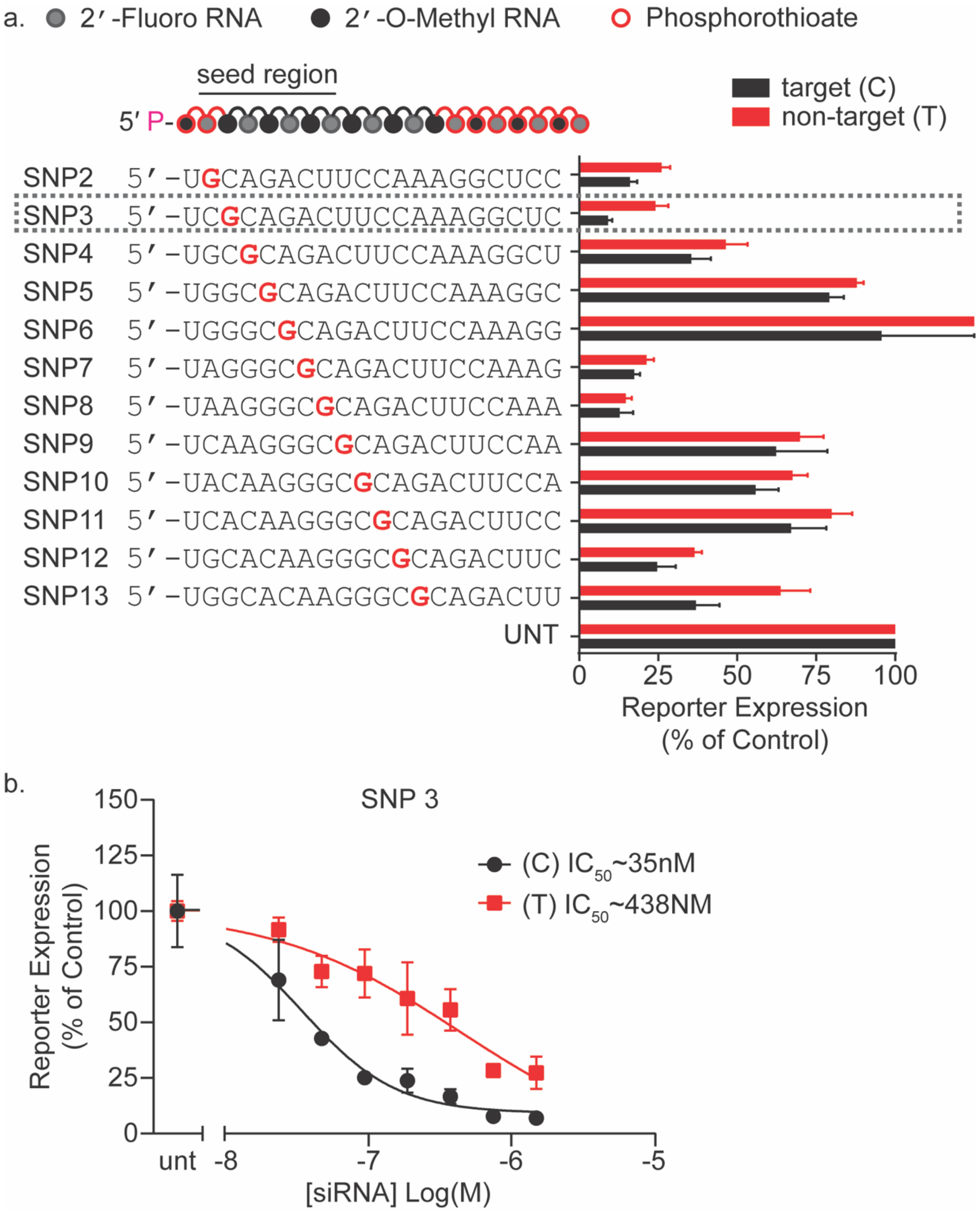
a.
2′-Fluoro RNA
2′-O-Methyl RNA
Phosphorothioate
seed region
5′ P-
target (C)
non-target (T)
SNP2 5′-UGCAGACUUCCAAAGGCUCC
SNP3 5′-UCGCAGACUUCCAAAGGCUC
SNP4 5′-UGCGCAGACUUCCAAAGGCU
SNP5 5′-UGGCGCAGACUUCCAAAGGC
SNP6 5′-UGGGCGCAGACUUCCAAAGG
SNP7 5′-UAGGGCGCAGACUUCCAAAG
SNP8 5′-UAAGGGCGCAGACUUCCAAA
SNP9 5′-UCAAGGGCGCAGACUUCCAA
SNP10 5′-UACAAGGGCGCAGACUUCCA
SNP11 5′-UCACAAGGGCGCAGACUUCC
SNP12 5′-UGCACAAGGGCGCAGACUUC
SNP13 5′-UGGCACAAGGGCGCAGACUU
UNT
0
25
50
75
100
Reporter Expression
(% of Control)
b.
SNP 3
150
125
100
75
50
25
0
Reporter Expression
(% of Control)
(C) IC50~35nM
(T) IC50~438NM
unt
-8
-7
-6
-5
[siRNA] Log(M)

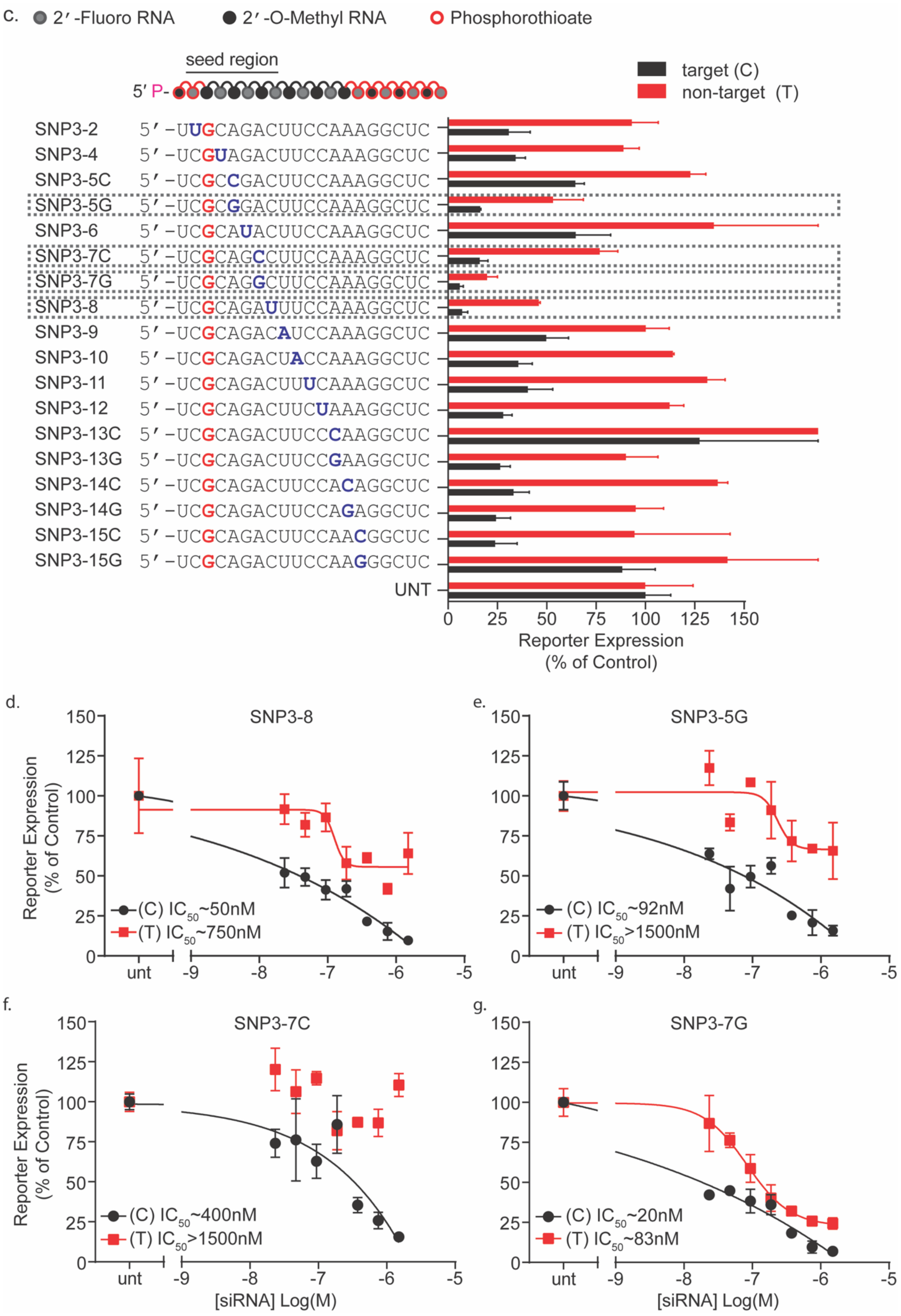
c.
2′-Fluoro RNA
2′-O-Methyl RNA
Phosphorothioate
seed region
5′ P-
target (C)
non-target (T)
SNP3-2 5′-UUGCAGACUUCCAAAGGCUC
SNP3-4 5′-UCGUAGACUUCCAAAGGCUC
SNP3-5C 5′-UCGCCGACUUCCAAAGGCUC
SNP3-5G 5′-UCGCGGACUUCCAAAGGCUC
SNP3-6 5′-UCGCAUACUUCCAAAGGCUC
SNP3-7C 5′-UCGCAGCCUUCCAAAGGCUC
SNP3-7G 5′-UCGCAGGCUUCCAAAGGCUC
SNP3-8 5′-UCGCAGAUUUCCAAAGGCUC
SNP3-9 5′-UCGCAGACAUCCAAAGGCUC
SNP3-10 5′-UCGCAGACUACCAAAGGCUC
SNP3-11 5′-UCGCAGACUUUCAAAGGCUC
SNP3-12 5′-UCGCAGACUUCUAAAGGCUC
SNP3-13C 5′-UCGCAGACUUCCCAAGGCUC
SNP3-13G 5′-UCGCAGACUUCCGAAGGCUC
SNP3-14C 5′-UCGCAGACUUCCACAGGCUC
SNP3-14G 5′-UCGCAGACUUCCAGAGGCUC
SNP3-15C 5′-UCGCAGACUUCCAACGGCUC
SNP3-15G 5′-UCGCAGACUUCCAAGGGCUC
UNT
0 25 50 75 100 125 150
Reporter Expression (% of Control)
d.
SNP3-8
(C) IC50~50nM
(T) IC50~750nM
e.
SNP3-5G
(C) IC50~92nM
(T) IC50>1500nM
f.
SNP3-7C
(C) IC50~400nM
(T) IC50>1500nM
g.
SNP3-7G
(C) IC50~20nM
(T) IC50~83nM
Reporter Expression (% of Control)
0 25 50 75 100 125 150
unt -9 -8 -7 -6 -5
[siRNA] Log(M)

***Supplementary Figure 5*. Screening method for sequence and mismatch selection can be used to produce effective SNP-selective siRNAs for a different SNP site. (a)** Results of primary screen finds optimal sequences for targeting SNP site rs362307 (highlighted in red). Compounds were tested using a dual-luciferase reporter assay system in HeLa cells. The (psiCheck) reporter plasmids contain a 40mer region of huntingtin, including the target SNP (C) (black), and non-target (U) isoform (red). Cells were treated for 72 hours at 1.5μM of siRNA. A panel of siRNA sequences were synthesized in a cholesterol-conjugated scaffold with phosphorothioate and alternating 2′-F and 2′-OMe backbone modifications. By walking the siRNA sequence around SNP site rs362307, we find multiple compounds with varying degrees of efficacy and discrimination. **(b)** Dose response of SNP3, which was selected for further screening of secondary mismatches in **(c)** Introduction of a secondary mismatch to positions 5 and 7 increases allelic discrimination. **(d-g)** A dose response of lead secondary-mismatch siRNAs validates an increase in discrimination based on the position of the mismatch. A C:U wobble mismatch at position 7 (SNP3-7C; **f**) results in a decrease in non-target activity when compared to a G:U mismatch at the same position (SNP3-7G; **g**).

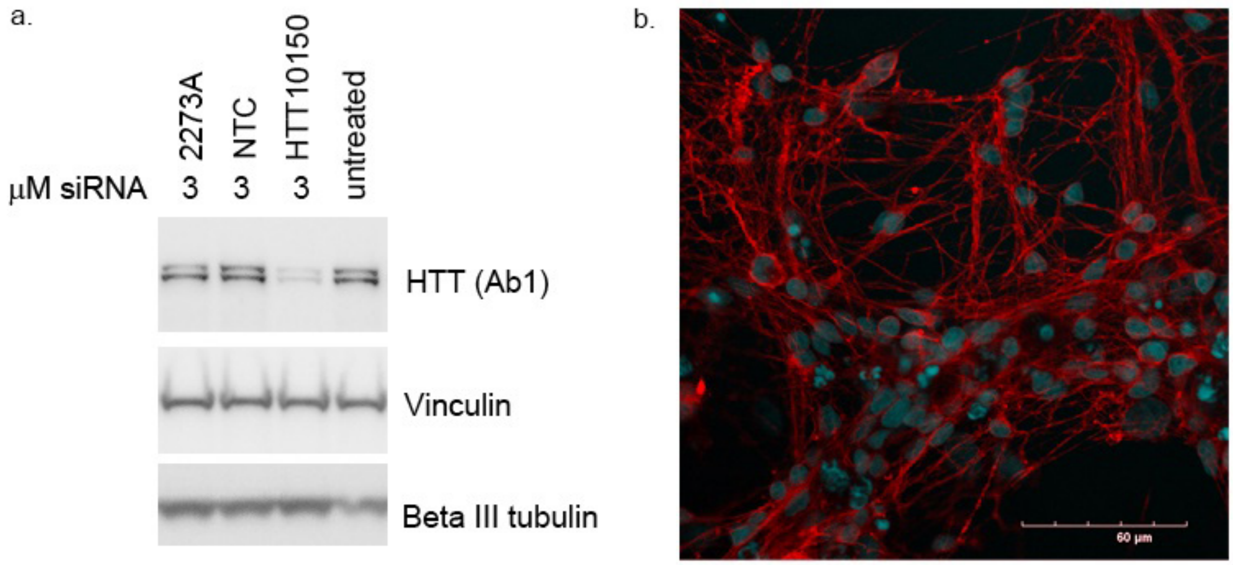


***Supplementary Figure 6.*** **Human HD109 neuron cultures show a high percentage of neurons. (a)** Western blot analysis of HD109 NSCs, which are heterozygous at SNP2273, showed lowering of mutant HTT (slower migrating band) compared to wild-type HTT (faster migrating band) with SNP2273A. For siRNA dosing regimen, see Methods. Equal protein (10 µg) were loaded per lane. Blots were probed with anti-HTT antibody Ab1 and vinculin as a housekeeping protein and the neuronal marker βIII tubulin. **(b)** Confocal immunofluorescent image of neuronal cultures stained for the neuronal marker βIII tubulin (Red) and the nuclear marker Hoechst (Blue) show high percentage of neurons in the cultures. Scale Bar=60mm ; 40X objective.

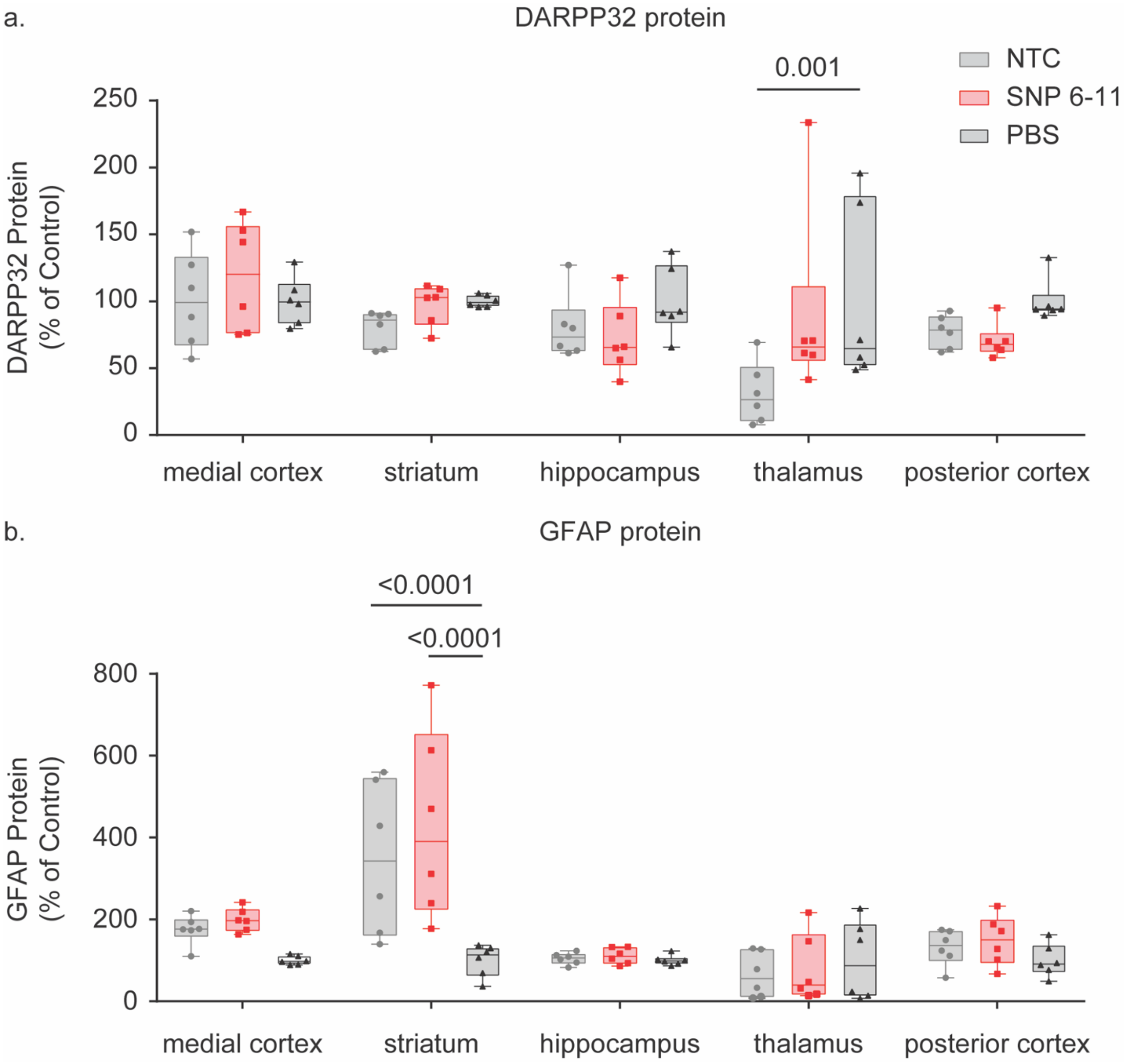


***Supplementary Figure 7.*** **Selective silencing of mutant *HTT* is well tolerated. (a)** SNP 6-11 siRNA has no impact on DARPP32 levels in treated mice although a decrease in DARPP-32 was observed in the thalamus of NTC-treated mice. **(b)** GFAP levels were also not affected by siRNA treatment, except in the striatum where an increase in GFAP was observed in both siRNA groups. A two-way ANOVA with multiple comparisons was performed for all protein analysis, comparing treatment groups to the PBS control for each brain region.

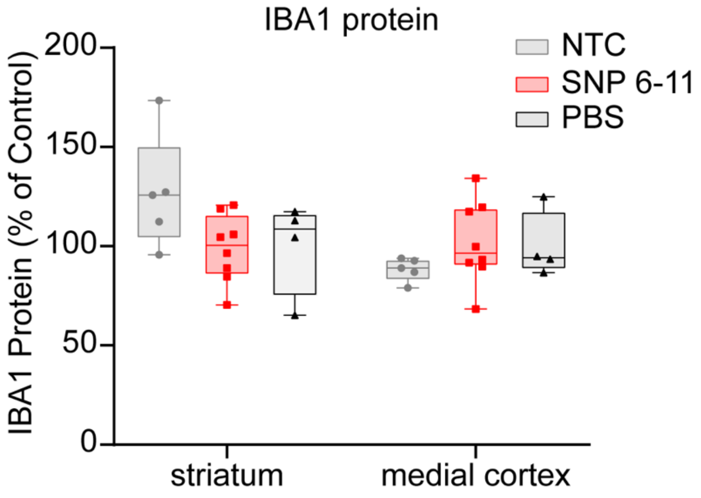


***Supplementary Figure 8.*** **SNP6-11 brain administration does not elicit microglial activation as indicated by lack of IBA1 elevation.** BAC-HD mice were injected with 225 μg of SNP6-11, NTC siRNAs and PBS. Levels of IBA1 expression in striatum and medial cortex, evaluated by automated western blot and normalized to GAPDH loading control. N=5-6, one-way ANOVA with Tukey multiple comparison correction.

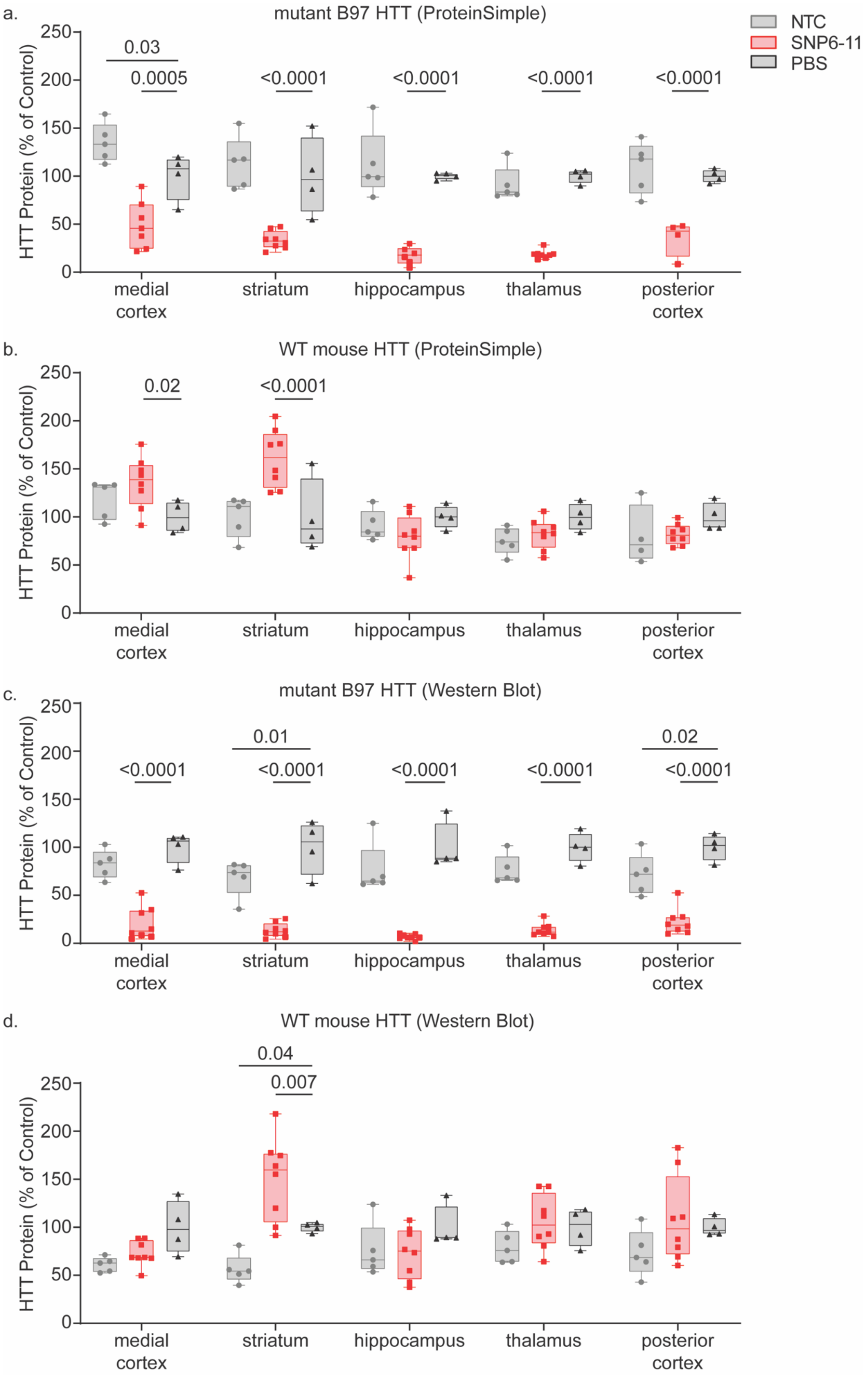
a.
mutant B97 HTT (ProteinSimple)
NTC
SNP6-11
PBS
HTT Protein (% of Control)
0.03
0.0005
<0.0001
<0.0001
<0.0001
<0.0001
medial cortex
striatum
hippocampus
thalamus
posterior cortex
b.
WT mouse HTT (ProteinSimple)
0.02
<0.0001
c.
mutant B97 HTT (Western Blot)
0.01
0.02
<0.0001
d.
WT mouse HTT (Western Blot)
0.04
0.007

***Supplementary Figure 9.*** **Increasing siRNA dose to 40nmols via ICV injection does not increase silencing of the HTT protein, or allelic discrimination. (a)** When treated with 450μg (20nmol, 10nmol/side) siRNA SNP6-11, selective silencing of mutant HTT protein (measured by WES Protein Simple) is achieved. There is an increase in wild-type HTT expression in the striatum and medial cortex compared to the controls, which is not seen at a 10nmol dose or in human cells. **(b)** Western blot shows that results are consistent among assays, with the exception of the medial cortex which shows an increase in WT HTT in the PBS control. A two-way ANOVA with multiple comparisons was performed for all protein analysis, comparing treatment groups to PBS control for each brain region.

***Table S1.*** complete list of all sequences of all siRNAs used in this study, with corresponding *in vitro* IC50 values of all compounds for which a dose response was performed.

| siRNA name | SNP position | additional mismatch position | antisense strand sequence | sense strand sequence | IC 50 (target) [nM] | IC50 (non-target) [nM] | Target SNP site | Related Figure (s) |
|---|---|---|---|---|---|---|---|---|
| SNP2-0 | 2 | 0 | P(mU)#(fU)#(mA)(fG)(mC)(fA)(mG)(fC)(mA)(fG)(mC)(fU)(mU)#(fC)#(mU)#(fC)#(mG)#(fU)#(mG)#(fG) | (fA)#(mG)#(fA)(mA)(fG)(mC)(fU)(mG)(fC)(mU)(fG)(mC)(fU)#(mA)#(fA)-TegChol | | | rs362273 | 1 |
| SNP3-0 | 3 | 0 | P(mU)#(fG)#(mU)(fA)(mG)(fC)(mA)(fG)(mC)(fA)(mG)(fC)(mU)#(fU)#(mC)#(fU)#(mC)#(fG)#(mU)#(fG) | (fG)#(mA)#(fA)(mG)(fC)(mU)(fG)(mC)(fU)(mG)(fC)(mU)(fA)#(mC)#(fA)-TegChol | | | rs362273 | 1 |
| SNP4-0 | 4 | 0 | P(mU)#(fU)#(mG)(fU)(mA)(fG)(mC)(fA)(mG)(fC)(mA)(fG)(mC)#(fU)#(mU)#(fC)#(mU)#(fC)#(mG)#(fU) | (fA)#(mA)#(fG)(mC)(fU)(mG)(fC)(mU)(fG)(mC)(fU)(mA)(fC)#(mA)#(fA)-TegChol | 38 | 803 | rs362273 | 1,S1 |
| SNP4-2 | 4 | 2 | P(mU)#(fA)#(mG)(fU)(mA)(fG)(mC)(fA)(mG)(fC)(mA)(fG)(mC)#(fU)#(mU)#(fC)#(mU)#(fC)#(mG)#(fU) | (fA)#(mA)#(fG)(mC)(fU)(mG)(fC)(mU)(fG)(mC)(fU)(mA)(fC)#(mA)#(fA)-TegChol | | | rs362273 | S4 |
| SNP4-3U | 4 | 3 | P(mU)#(fU)#(mU)(fU)(mA)(fG)(mC)(fA)(mG)(fC)(mA)(fG)(mC)#(fU)#(mU)#(fC)#(mU)#(fC)#(mG)#(fU) | (fA)#(mA)#(fG)(mC)(fU)(mG)(fC)(mU)(fG)(mC)(fU)(mA)(fC)#(mA)#(fA)-TegChol | | | rs362273 | S4 |
| SNP4-3A | 4 | 3 | P(mU)#(fU)#(mA)(fU)(mA)(fG)(mC)(fA)(mG)(fC)(mA)(fG)(mC)#(fU)#(mU)#(fC)#(mU)#(fC)#(mG)#(fU) | (fA)#(mA)#(fG)(mC)(fU)(mG)(fC)(mU)(fG)(mC)(fU)(mA)(fC)#(mA)#(fA)-TegChol | | | rs362273 | S4 |
| SNP4-5C | 4 | 5 | P(mU)#(fU)#(mG)(fU)(mC)(fG)(mC)(fA)(mG)(fC)(mA)(fG)(mC)#(fU)#(mU)#(fC)#(mU)#(fC)#(mG)#(fU) | (fA)#(mA)#(fG)(mC)(fU)(mG)(fC)(mU)(fG)(mC)(fU)(mA)(fC)#(mA)#(fA)-TegChol | | | rs362273 | S4 |
| SNP4-5G | 4 | 5 | P(mU)#(fU)#(mG)(fU)(mG)(fG)(mC)(fA)(mG)(fC)(mA)(fG)(mC)#(fU)#(mU)#(fC)#(mU)#(fC)#(mG)#(fU) | (fA)#(mA)#(fG)(mC)(fU)(mG)(fC)(mU)(fG)(mC)(fU)(mA)(fC)#(mA)#(fA)-TegChol | | | rs362273 | S4 |
| SNP4-6 | 4 | 6 | P(mU)#(fU)#(mG)(fU)(mA)(fU)(mC)(fA)(mG)(fC)(mA)(fG)(mC)#(fU)#(mU)#(fC)#(mU)#(fC)#(mG)#(fU) | (fA)#(mA)#(fG)(mC)(fU)(mG)(fC)(mU)(fG)(mC)(fU)(mA)(fC)#(mA)#(fA)-TegChol | | | rs362273 | S4 |
| SNP4-7 | 4 | 7 | P(mU)#(fU)#(mG)(fU)(mA)(fG)(mU)(fA)(mG)(fC)(mA)(fG)(mC)#(fU)#(mU)#(fC)#(mU)#(fC)#(mG)#(fU) | (fA)#(mA)#(fG)(mC)(fU)(mG)(fC)(mU)(fG)(mC)(fU)(mA)(fC)#(mA)#(fA)-TegChol | 17 | >5000 | rs362273 | S4 |
| SNP4-8 | 4 | 8 | P(mU)#(fU)#(mG)(fU)(mA)(fG)(mC)(fU)(mG)(fC)(mA)(fG)(mC)#(fU)#(mU)#(fC)#(mU)#(fC)#(mG)#(fU) | (fA)#(mA)#(fG)(mC)(fU)(mG)(fC)(mU)(fG)(mC)(fU)(mA)(fC)#(mA)#(fA)-TegChol | 24 | 220 | rs362273 | S4 |
| SNP4-9 | 4 | 9 | P(mU)#(fU)#(mG)(fU)(mA)(fG)(mC)(fA)(mU)(fC)(mA)(fG)(mC)#(fU)#(mU)#(fC)#(mU)#(fC)#(mG)#(fU) | (fA)#(mA)#(fG)(mC)(fU)(mG)(fC)(mU)(fG)(mC)(fU)(mA)(fC)#(mA)#(fA)-TegChol | | | rs362273 | S4 |
| SNP4-10 | 4 | 10 | P(mU)#(fU)#(mG)(fU)(mA)(fG)(mC)(fA)(mG)(fU)(mA)(fG)(mC)#(fU)#(mU)#(fC)#(mU)#(fC)#(mG)#(fU) | (fA)#(mA)#(fG)(mC)(fU)(mG)(fC)(mU)(fG)(mC)(fU)(mA)(fC)#(mA)#(fA)-TegChol | | | rs362273 | S4 |
| SNP4-11 | 4 | 11 | P(mU)#(fU)#(mG)(fU)(mA)(fG)(mC)(fA)(mG)(fC)(mU)(fG)(mC)#(fU)#(mU)#(fC)#(mU)#(fC)#(mG)#(fU) | (fA)#(mA)#(fG)(mC)(fU)(mG)(fC)(mU)(fG)(mC)(fU)(mA)(fC)#(mA)#(fA)-TegChol | | | rs362273 | S4 |
| SNP4-12 | 4 | 12 | P(mU)#(fU)#(mG)(fU)(mA)(fG)(mC)(fA)(mG)(fC)(mA)(fU)(mC)#(fU)#(mU)#(fC)#(mU)#(fC)#(mG)#(fU) | (fA)#(mA)#(fG)(mC)(fU)(mG)(fC)(mU)(fG)(mC)(fU)(mA)(fC)#(mA)#(fA)-TegChol | | | rs362273 | S4 |
| SNP4-13 | 4 | 13 | P(mU)#(fU)#(mG)(fU)(mA)(fG)(mC)(fA)(mG)(fC)(mA)(fG)(mU)#(fU)#(mU)#(fC)#(mU)#(fC)#(mG)#(fU) | (fA)#(mA)#(fG)(mC)(fU)(mG)(fC)(mU)(fG)(mC)(fU)(mA)(fC)#(mA)#(fA)-TegChol | | | rs362273 | S4 |
| SNP4-14 | 4 | 14 | P(mU)#(fU)#(mG)(fU)(mA)(fG)(mC)(fA)(mG)(fC)(mA)(fG)(mC)#(fA)#(mU)#(fC)#(mU)#(fC)#(mG)#(fU) | (fA)#(mA)#(fG)(mC)(fU)(mG)(fC)(mU)(fG)(mC)(fU)(mA)(fC)#(mA)#(fA)-TegChol | | | rs362273 | S4 |
| SNP4-15 | 4 | 15 | P(mU)#(fU)#(mG)(fU)(mA)(fG)(mC)(fA)(mG)(fC)(mA)(fG)(mC)#(fU)#(mA)#(fC)#(mU)#(fC)#(mG)#(fU) | (fA)#(mA)#(fG)(mC)(fU)(mG)(fC)(mU)(fG)(mC)(fU)(mA)(fC)#(mA)#(fA)-TegChol | | | rs362273 | S4 |
| SNP4-16 | 4 | 16 | P(mU)#(fU)#(mG)(fU)(mA)(fG)(mC)(fA)(mG)(fC)(mA)(fG)(mC)#(fU)#(mU)#(fU)#(mU)#(fC)#(mG)#(fU) | (fA)#(mA)#(fG)(mC)(fU)(mG)(fC)(mU)(fG)(mC)(fU)(mA)(fC)#(mA)#(fA)-TegChol | | | rs362273 | S4 |
| SNP5-0 | 5 | 0 | P(mU)#(fC)#(mU)(fG)(mU)(fA)(mG)(fC)(mA)(fG)(mC)(fA)(mG)#(fC)#(mU)#(fU)#(mC)#(fU)#(mC)#(fG) | (fA)#(mG)#(fC)(mU)(fG)(mC)(fU)(mG)(fC)(mU)(fA)(mC)(fA)#(mG)#(fA)-TegChol | | | rs362273 | 1 |
| SNP6-0 | 6 | 0 | P(mU)#(fU)#(mC)(fU)(mG)(fU)(mA)(fG)(mC)(fA)(mG)(fC)(mA)#(fG)#(mC)#(fU)#(mU)#(fC)#(mU)#(fC) | (fG)#(mC)#(fU)(mG)(fC)(mU)(fG)(mC)(fU)(mA)(fC)(mA)(fG)#(mA)#(fA)-TegChol | 24 | 174 | rs362273 | 1,S1,2 |
| SNP6-2 | 6 | 2 | P(mU)#(fA)#(mC)(fU)(mG)(fU)(mA)(fG)(mC)(fA)(mG)(fC)(mA)#(fG)#(mC)#(fU)#(mU)#(fC)#(mU)#(fC) | (fG)#(mC)#(fU)(mG)(fC)(mU)(fG)(mC)(fU)(mA)(fC)(mA)(fG)#(mA)#(fA)-TegChol | | | rs362273 | 2 |
| SNP6-3 | 6 | 3 | P(mU)#(fU)#(mU)(fU)(mG)(fU)(mA)(fG)(mC)(fA)(mG)(fC)(mA)#(fG)#(mC)#(fU)#(mU)#(fC)#(mU)#(fC) | (fG)#(mC)#(fU)(mG)(fC)(mU)(fG)(mC)(fU)(mA)(fC)(mA)(fG)#(mA)#(fA)-TegChol | | | rs362273 | 2 |
| SNP6-4 | 6 | 4 | P(mU)#(fU)#(mC)(fA)(mG)(fU)(mA)(fG)(mC)(fA)(mG)(fC)(mA)#(fG)#(mC)#(fU)#(mU)#(fC)#(mU)#(fC) | (fG)#(mC)#(fU)(mG)(fC)(mU)(fG)(mC)(fU)(mA)(fC)(mA)(fG)#(mA)#(fA)-TegChol | | | rs362273 | 2 |
| SNP6-5U | 6 | 5 | P(mU)#(fU)#(mC)(fU)(mU)(fU)(mA)(fG)(mC)(fA)(mG)(fC)(mA)#(fG)#(mC)#(fU)#(mU)#(fC)#(mU)#(fC) | (fG)#(mC)#(fU)(mG)(fC)(mU)(fG)(mC)(fU)(mA)(fC)(mA)(fG)#(mA)#(fA)-TegChol | | | rs362273 | 2 |
| SNP6-5A | 6 | 5 | P(mU)#(fU)#(mC)(fU)(mA)(fU)(mA)(fG)(mC)(fA)(mG)(fC)(mA)#(fG)#(mC)#(fU)#(mU)#(fC)#(mU)#(fC) | (fG)#(mC)#(fU)(mG)(fC)(mU)(fG)(mC)(fU)(mA)(fC)(mA)(fG)#(mA)#(fA)-TegChol | | | rs362273 | 2 |
| SNP6-7C | 6 | 7 | P(mU)#(fU)#(mC)(fU)(mG)(fU)(mC)(fG)(mC)(fA)(mG)(fC)(mA)#(fG)#(mC)#(fU)#(mU)#(fC)#(mU)#(fC) | (fG)#(mC)#(fU)(mG)(fC)(mU)(fG)(mC)(fU)(mA)(fC)(mA)(fG)#(mA)#(fA)-TegChol | | | rs362273 | 2 |
| SNP6-7G | 6 | 7 | P(mU)#(fU)#(mC)(fU)(mG)(fU)(mG)(fG)(mC)(fA)(mG)(fC)(mA)#(fG)#(mC)#(fU)#(mU)#(fC)#(mU)#(fC) | (fG)#(mC)#(fU)(mG)(fC)(mU)(fG)(mC)(fU)(mA)(fC)(mA)(fG)#(mA)#(fA)-TegChol | | | rs362273 | 2 |
| SNP6-8 | 6 | 8 | P(mU)#(fU)#(mC)(fU)(mG)(fU)(mA)(fU)(mC)(fA)(mG)(fC)(mA)#(fG)#(mC)#(fU)#(mU)#(fC)#(mU)#(fC) | (fG)#(mC)#(fU)(mG)(fC)(mU)(fG)(mC)(fU)(mA)(fC)(mA)(fG)#(mA)#(fA)-TegChol | | | rs362273 | 2 |
| SNP6-9 | 6 | 9 | P(mU)#(fU)#(mC)(fU)(mG)(fU)(mA)(fG)(mU)(fA)(mG)(fC)(mA)#(fG)#(mC)#(fU)#(mU)#(fC)#(mU)#(fC) | (fG)#(mC)#(fU)(mG)(fC)(mU)(fG)(mC)(fU)(mA)(fC)(mA)(fG)#(mA)#(fA)-TegChol | | | rs362273 | 2 |
| SNP6-10 | 6 | 10 | P(mU)#(fU)#(mC)(fU)(mG)(fU)(mA)(fG)(mC)(fU)(mG)(fC)(mA)#(fG)#(mC)#(fU)#(mU)#(fC)#(mU)#(fC) | (fG)#(mC)#(fU)(mG)(fC)(mU)(fG)(mC)(fU)(mA)(fC)(mA)(fG)#(mA)#(fA)-TegChol | | | rs362273 | 2 |
| SNP6-11 (G) | 6 | 11 | P(mU)#(fU)#(mC)(fU)(mG)(fC)(mA)(fG)(mC)(fA)(mU)(fC)(mA)#(fG)#(mC)#(fU)#(mU)#(fC)#(mU)#(fC) | (fG)#(mC)#(fU)(mG)(fC)(mU)(fG)(mC)(fU)(mG)(fC)(mA)(fG)#(mA)#(fA)-TegChol | | | rs362273 | 2,4,S3 |
| SNP6-11 (A) | 6 | 11 | P(mU)#(fU)#(mC)(fU)(mG)(fU)(mA)(fG)(mC)(fA)(mU)(fC)(mA)#(fG)#(mC)#(fU)#(mU)#(fC)#(mU)#(fC) | (fG)#(mC)#(fU)(mG)(fC)(mU)(fG)(mC)(fU)(mA)(fC)(mA)(fG)#(mA)#(fA)-TegChol | 28 | >4500 | rs362273 | 2,3,4,S3 |
| SNP6-12 | 6 | 12 | P(mU)#(fU)#(mC)(fU)(mG)(fU)(mA)(fG)(mC)(fA)(mG)(fU)(mA)#(fG)#(mC)#(fU)#(mU)#(fC)#(mU)#(fC) | (fG)#(mC)#(fU)(mG)(fC)(mU)(fG)(mC)(fU)(mA)(fC)(mA)(fG)#(mA)#(fA)-TegChol | 24 | >1000 | rs362273 | 2 |
| SNP6-13 | 6 | 13 | P(mU)#(fU)#(mC)(fU)(mG)(fU)(mA)(fG)(mC)(fA)(mG)(fC)(mU)#(fG)#(mC)#(fU)#(mU)#(fC)#(mU)#(fC) | (fG)#(mC)#(fU)(mG)(fC)(mU)(fG)(mC)(fU)(mA)(fC)(mA)(fG)#(mA)#(fA)-TegChol | | | rs362273 | 2 |
| SNP6-14 | 6 | 14 | P(mU)#(fU)#(mC)(fU)(mG)(fU)(mA)(fG)(mC)(fA)(mG)(fC)(mA)#(fU)#(mC)#(fU)#(mU)#(fC)#(mU)#(fC) | (fG)#(mC)#(fU)(mG)(fC)(mU)(fG)(mC)(fU)(mA)(fC)(mA)(fG)#(mA)#(fA)-TegChol | 30 | >2000 | rs362273 | 2 |
| SNP6-15 | 6 | 15 | P(mU)#(fU)#(mC)(fU)(mG)(fU)(mA)(fG)(mC)(fA)(mG)(fC)(mA)#(fG)#(mU)#(fU)#(mU)#(fC)#(mU)#(fC) | (fG)#(mC)#(fU)(mG)(fC)(mU)(fG)(mC)(fU)(mA)(fC)(mA)(fG)#(mA)#(fA)-TegChol | | | rs362273 | 2 |
| SNP6-16 | 6 | 16 | P(mU)#(fU)#(mC)(fU)(mG)(fU)(mA)(fG)(mC)(fA)(mG)(fC)(mA)#(fG)#(mC)#(fA)#(mU)#(fC)#(mU)#(fC) | (fG)#(mC)#(fU)(mG)(fC)(mU)(fG)(mC)(fU)(mA)(fC)(mA)(fG)#(mA)#(fA)-TegChol | 18 | 333 | rs362273 | 2 |
| SNP7-0 | 7 | 0 | P(mU)#(fA)#(mU)(fC)(mU)(fG)(mU)(fA)(mG)(fC)(mA)(fG)(mC)#(fA)#(mG)#(fC)#(mU)#(fU)#(mC)#(fU) | (fC)#(mU)#(fG)(mC)(fU)(mG)(fC)(mU)(fA)(mC)(fA)(mG)(fA)#(mU)#(fA)-TegChol | | | rs362273 | 1 |
| SNP8-0 | 8 | 0 | P(mU)#(fG)#(mA)(fU)(mC)(fU)(mG)(fU)(mA)(fG)(mC)(fA)(mG)#(fC)#(mA)#(fG)#(mC)#(fU)#(mU)#(fC) | (fU)#(mG)#(fC)(mU)(fG)(mC)(fU)(mA)(fC)(mA)(fG)(mA)(fU)#(mC)#(fA)-TegChol | | | rs362273 | 1 |
| SNP9-0 | 9 | 0 | P(mU)#(fU)#(mG)(fA)(mU)(fC)(mU)(fG)(mU)(fA)(mG)(fC)(mA)#(fG)#(mC)#(fA)#(mG)#(fC)#(mU)#(fU) | (fG)#(mC)#(fU)(mG)(fC)(mU)(fA)(mC)(fA)(mG)(fA)(mU)(fC)#(mA)#(fA)-TegChol | | | rs362273 | 1 |
| SNP10-0 | 10 | 0 | P(mU)#(fU)#(mU)(fG)(mA)(fU)(mC)(fU)(mG)(fU)(mA)(fG)(mC)#(fA)#(mG)#(fC)#(mA)#(fG)#(mC)#(fU) | (fC)#(mU)#(fG)(mC)(fU)(mA)(fC)(mA)(fG)(mA)(fU)(mC)(fA)#(mA)#(fA)-TegChol | | | rs362273 | 1 |
| SNP11-0 | 11 | 0 | P(mU)#(fG)#(mU)(fU)(mG)(fA)(mU)(fC)(mU)(fG)(mU)(fA)(mG)#(fC)#(mA)#(fG)#(mC)#(fA)#(mG)#(fC) | (fU)#(mG)#(fC)(mU)(fA)(mC)(fA)(mG)(fA)(mU)(fC)(mA)(fA)#(mC)#(fA)-TegChol | | | rs362273 | 1 |
| SNP12-0 | 12 | 0 | P(mU)#(fG)#(mG)(fU)(mU)(fG)(mA)(fU)(mC)(fU)(mG)(fU)(mA)#(fG)#(mC)#(fA)#(mG)#(fC)#(mA)#(fG) | (fG)#(mC)#(fU)(mA)(fC)(mA)(fG)(mA)(fU)(mC)(fA)(mA)(fC)#(mC)#(fA)-TegChol | | | rs362273 | 1 |
| SNP13-0 | 13 | 0 | P(mU)#(fG)#(mG)(fG)(mU)(fU)(mG)(fA)(mU)(fC)(mU)(fG)(mU)#(fA)#(mG)#(fC)#(mA)#(fG)#(mC)#(fA) | (fC)#(mU)#(fA)(mC)(fA)(mG)(fA)(mU)(fC)(mA)(fA)(mC)(fC)#(mC)#(fA)-TegChol | | | rs362273 | 1 |
| SNP 6-f11 | 6 | 11 | P(mU)#(fU)#(mC)(fU)(mG)(fU)(mA)(fG)(mC)(fA)(fU)(fC)(mA)#(fG)#(mC)#(fU)#(mU)#(fC)#(mU)#(fC) | (fG)#(mC)#(fU)(mG)(mC)(mU)(fG)(mC)(fU)(mA)(fC)(mA)(fG)#(mA)#(fA)-TegChol | | | rs362273 | 3 |
| SNP f6-f11 | 6 | 11 | P(mU)#(fU)#(mC)(fU)(fG)(fU)(fA)(fG)(mC)(fA)(fU)(fC)(mA)#(fG)#(mC)#(fU)#(mU)#(fC)#(mU)#(fC) | (fG)#(mC)#(fU)(mG)(mC)(mU)(fG)(mC)(mU)(mA)(mC)(mA)(fG)#(mA)#(fA)-TegChol | | | rs362273 | 3 |
| SNP f6-m11 | 6 | 11 | P(mU)#(fU)#(mC)(fU)(fG)(fU)(fA)(fG)(mC)(mA)(mU)(mC)(mA)#(fG)#(mC)#(fU)#(mU)#(fC)#(mU)#(fC) | (fG)#(mC)#(fU)(fG)(fC)(fU)(fG)(mC)(mU)(mA)(mC)(mA)(fG)#(mA)#(fA)-TegChol | | | rs362273 | 3 |
| SNP m6-f11 | 6 | 11 | P(mU)#(fU)#(mC)(fU)(mG)(mU)(mA)(fG)(mC)(fA)(fU)(fC)(mA)#(fG)#(mC)#(fU)#(mU)#(fC)#(mU)#(fC) | (fG)#(mC)#(fU)(mG)(mC)(mU)(fG)(mC)(fU)(fA)(fC)(mA)(fG)#(mA)#(fA)-TegChol | | | rs362273 | 3 |
| SNP m6-m11 | 6 | 11 | P(mU)#(fU)#(mC)(fU)(mG)(mU)(mA)(fG)(mC)(mA)(mU)(mC)(mA)#(fG)#(mC)#(fU)#(mU)#(fC)#(mU)#(fC) | (fG)#(mC)#(fU)(fG)(fC)(fU)(fG)(mC)(fU)(fA)(fC)(mA)(fG)#(mA)#(fA)-TegChol | | | rs362273 | 3 |
| SNP 6-m11 | 6 | 11 | P(mU)#(fU)#(mC)(fU)(mG)(fU)(mA)(fG)(mC)(mA)(mU)(mC)(mA)#(fG)#(mC)#(fU)#(mU)#(fC)#(mU)#(fC) | (fG)#(mC)#(fU)(fG)(fC)(fU)(fG)(mC)(fU)(mA)(fC)(mA)(fG)#(mA)#(fA)-TegChol | | | rs362273 | 3 |

| | | | | | | | | |
|---|---|---|---|---|---|---|---|---|
| SNP fm6-11 | 6 | 11 | P(mU)#(fU)#(mC)(fU)(fG)(mU)(fA)(fG)(mC)(fA)(mU)(fC)(mA)#(fG)#(mC)#(fU)#(mU)#(fC)#(mU)#(fC) | (fG)#(mC)#(fU)(mG)(fC)(mU)(fG)(mC)(mU)(fA)(mC)(mA)(fG)#(mA)#(fA)-TegChol | | | rs362273 | 3 |
| SNP f6-11 | 6 | 11 | P(mU)#(fU)#(mC)(fU)(fG)(fU)(fA)(fG)(mC)(fA)(mU)(fC)(mA)#(fG)#(mC)#(fU)#(mU)#(fC)#(mU)#(fC) | (fG)#(mC)#(fU)(mG)(fC)(mU)(fG)(mC)(mU)(mA)(mC)(mA)(fG)#(mA)#(fA)-TegChol | | | rs362273 | 3 |
| SNP m6-11 | 6 | 11 | P(mU)#(fU)#(mC)(fU)(mG)(mU)(mA)(fG)(mC)(fA)(mU)(fC)(mA)#(fG)#(mC)#(fU)#(mU)#(fC)#(mU)#(fC) | (fG)#(mC)#(fU)(mG)(fC)(mU)(fG)(mC)(fU)(fA)(fC)(mA)(fG)#(mA)#(fA)-TegChol | | | rs362273 | 3 |
| SNP2 | 2 | 0 | P(mU)#(fG)#(mC)(fA)(mG)(fA)(mC)(fU)(mU)(fC)(mC)(fA)(mA)#(fA)#(mG)#(fG)#(mC)#(fU)#(mC)#(fC) | (fC)#(mU)#(fU)(mU)(fG)(mG)(fA)(mA)(fG)(mU)(fC)(mU)(fG)#(mC)#(fA)-TegChol | | | rs362307 | S5 |
| SNP3 | 3 | 0 | P(mU)#(fC)#(mG)(fC)(mA)(fG)(mA)(fC)(mU)(fU)(mC)(fC)(mA)#(fA)#(mA)#(fG)#(mG)#(fC)#(mU)#(fC) | (fU)#(mU)#(fU)(mG)(fG)(mA)(fA)(mG)(fU)(mC)(fU)(mG)(fC)#(mG)#(fA)-TegChol | 35 | 438 | rs362307 | S5 |
| SNP4 | 4 | 0 | P(mU)#(fG)#(mC)(fG)(mC)(fA)(mG)(fA)(mC)(fU)(mU)(fC)(mC)#(fA)#(mA)#(fA)#(mG)#(fG)#(mC)#(fU) | (fU)#(mU)#(fG)(mG)(fA)(mA)(fG)(mU)(fC)(mU)(fG)(mC)(fG)#(mC)#(fA)-TegChol | | | rs362307 | S5 |
| SNP5 | 5 | 0 | P(mU)#(fG)#(mG)(fC)(mG)(fC)(mA)(fG)(mA)(fC)(mU)(fU)(mC)#(fC)#(mA)#(fA)#(mA)#(fG)#(mG)#(fC) | (fU)#(mG)#(fG)(mA)(fA)(mG)(fU)(mC)(fU)(mG)(fC)(mG)(fC)#(mC)#(fA)-TegChol | | | rs362307 | S5 |
| SNP6 | 6 | 0 | P(mU)#(fG)#(mG)(fG)(mC)(fG)(mC)(fA)(mG)(fA)(mC)(fU)(mU)#(fC)#(mC)#(fA)#(mA)#(fA)#(mG)#(fG) | (fG)#(mG)#(fA)(mA)(fG)(mU)(fC)(mU)(fG)(mC)(fG)(mC)(fC)#(mC)#(fA)-TegChol | | | rs362307 | S5 |
| SNP7 | 7 | 0 | P(mU)#(fA)#(mG)(fG)(mG)(fC)(mG)(fC)(mA)(fG)(mA)(fC)(mU)#(fU)#(mC)#(fC)#(mA)#(fA)#(mA)#(fG) | (fG)#(mA)#(fA)(mG)(fU)(mC)(fU)(mG)(fC)(mG)(fC)(mC)(fC)#(mU)#(fA)-TegChol | | | rs362307 | S5 |
| SNP8 | 8 | 0 | P(mU)#(fA)#(mA)(fG)(mG)(fG)(mC)(fG)(mC)(fA)(mG)(fA)(mC)#(fU)#(mU)#(fC)#(mC)#(fA)#(mA)#(fA) | (fA)#(mA)#(fG)(mU)(fC)(mU)(fG)(mC)(fG)(mC)(fC)(mC)(fU)#(mU)#(fA)-TegChol | | | rs362307 | S5 |
| SNP9 | 9 | 0 | P(mU)#(fC)#(mA)(fA)(mG)(fG)(mG)(fC)(mG)(fC)(mA)(fG)(mA)#(fC)#(mU)#(fU)#(mC)#(fC)#(mA)#(fA) | (fA)#(mG)#(fU)(mC)(fU)(mG)(fC)(mG)(fC)(mC)(fC)(mU)(fU)#(mG)#(fA)-TegChol | | | rs362307 | S5 |
| SNP10 | 10 | 0 | P(mU)#(fA)#(mC)(fA)(mA)(fG)(mG)(fG)(mC)(fG)(mC)(fA)(mG)#(fA)#(mC)#(fU)#(mU)#(fC)#(mC)#(fA) | (fG)#(mU)#(fC)(mU)(fG)(mC)(fG)(mC)(fC)(mC)(fU)(mU)(fG)#(mU)#(fA)-TegChol | | | rs362307 | S5 |
| SNP11 | 11 | 0 | P(mU)#(fC)#(mA)(fC)(mA)(fA)(mG)(fG)(mG)(fC)(mG)(fC)(mA)#(fG)#(mA)#(fC)#(mU)#(fU)#(mC)#(fC) | (fU)#(mC)#(fU)(mG)(fC)(mG)(fC)(mC)(fC)(mU)(fU)(mG)(fU)#(mG)#(fA)-TegChol | | | rs362307 | S5 |
| SNP12 | 12 | 0 | P(mU)#(fG)#(mC)(fA)(mC)(fA)(mA)(fG)(mG)(fG)(mC)(fG)(mC)#(fA)#(mG)#(fA)#(mC)#(fU)#(mU)#(fC) | (fC)#(mU)#(fG)(mC)(fG)(mC)(fC)(mC)(fU)(mU)(fG)(mU)(fG)#(mC)#(fA)-TegChol | | | rs362307 | S5 |
| SNP13 | 13 | 0 | P(mU)#(fG)#(mG)(fC)(mA)(fC)(mA)(fA)(mG)(fG)(mG)(fC)(mG)#(fC)#(mA)#(fG)#(mA)#(fC)#(mU)#(fU) | (fU)#(mG)#(fC)(mG)(fC)(mC)(fC)(mU)(fU)(mG)(fU)(mG)(fC)#(mC)#(fA)-TegChol | | | rs362307 | S5 |
| SNP 3-2 | 3 | 2 | P(mU)#(fU)#(mG)(fC)(mA)(fG)(mA)(fC)(mU)(fU)(mC)(fC)(mA)#(fA)#(mA)#(fG)#(mG)#(fC)#(mU)#(fC) | (fU)#(mU)#(fU)(mG)(fG)(mA)(fA)(mG)(fU)(mC)(fU)(mG)(fC)#(mA)#(fA)-TegChol | | | rs362307 | S5 |
| SNP 3-4 | 3 | 4 | P(mU)#(fC)#(mG)(fU)(mA)(fG)(mA)(fC)(mU)(fU)(mC)(fC)(mA)#(fA)#(mA)#(fG)#(mG)#(fC)#(mU)#(fC) | (fU)#(mU)#(fU)(mG)(fG)(mA)(fA)(mG)(fU)(mC)(fU)(mA)(fC)#(mG)#(fA)-TegChol | | | rs362307 | S5 |
| SNP 3-5C | 3 | 5 | P(mU)#(fC)#(mG)(fC)(mC)(fG)(mA)(fC)(mU)(fU)(mC)(fC)(mA)#(fA)#(mA)#(fG)#(mG)#(fC)#(mU)#(fC) | (fU)#(mU)#(fU)(mG)(fG)(mA)(fA)(mG)(fU)(mC)(fG)(mG)(fC)#(mG)#(fA)-TegChol | 92 | >1500 | rs362307 | S5 |
| SNP 3-5G | 3 | 5 | P(mU)#(fC)#(mG)(fC)(mG)(fG)(mA)(fC)(mU)(fU)(mC)(fC)(mA)#(fA)#(mA)#(fG)#(mG)#(fC)#(mU)#(fC) | (fU)#(mU)#(fU)(mG)(fG)(mA)(fA)(mG)(fU)(mC)(fC)(mG)(fC)#(mG)#(fA)-TegChol | | | rs362307 | S5 |
| SNP 3-6 | 3 | 6 | P(mU)#(fC)#(mG)(fC)(mA)(fU)(mA)(fC)(mU)(fU)(mC)(fC)(mA)#(fA)#(mA)#(fG)#(mG)#(fC)#(mU)#(fC) | (fU)#(mU)#(fU)(mG)(fG)(mA)(fA)(mG)(fU)(mA)(fU)(mG)(fC)#(mG)#(fA)-TegChol | | | rs362307 | S5 |
| SNP 3-7C | 3 | 7 | P(mU)#(fC)#(mG)(fC)(mA)(fG)(mC)(fC)(mU)(fU)(mC)(fC)(mA)#(fA)#(mA)#(fG)#(mG)#(fC)#(mU)#(fC) | (fU)#(mU)#(fU)(mG)(fG)(mA)(fA)(mG)(fG)(mC)(fU)(mG)(fC)#(mG)#(fA)-TegChol | 400 | >1500 | rs362307 | S5 |
| SNP 3-7G | 3 | 7 | P(mU)#(fC)#(mG)(fC)(mA)(fG)(mG)(fC)(mU)(fU)(mC)(fC)(mA)#(fA)#(mA)#(fG)#(mG)#(fC)#(mU)#(fC) | (fU)#(mU)#(fU)(mG)(fG)(mA)(fA)(mG)(fC)(mC)(fU)(mG)(fC)#(mG)#(fA)-TegChol | 20 | 83 | rs362307 | S5 |
| SNP 3-8 | 3 | 8 | P(mU)#(fC)#(mG)(fC)(mA)(fG)(mA)(fU)(mU)(fU)(mC)(fC)(mA)#(fA)#(mA)#(fG)#(mG)#(fC)#(mU)#(fC) | (fU)#(mU)#(fU)(mG)(fG)(mA)(fA)(mA)(fU)(mC)(fU)(mG)(fC)#(mG)#(fA)-TegChol | 50 | 750 | rs362307 | S5 |
| SNP 3-9 | 3 | 9 | P(mU)#(fC)#(mG)(fC)(mA)(fG)(mA)(fC)(mA)(fU)(mC)(fC)(mA)#(fA)#(mA)#(fG)#(mG)#(fC)#(mU)#(fC) | (fU)#(mU)#(fU)(mG)(fG)(mA)(fU)(mG)(fU)(mC)(fU)(mG)(fC)#(mG)#(fA)-TegChol | | | rs362307 | S5 |
| SNP 3-10 | 3 | 10 | P(mU)#(fC)#(mG)(fC)(mA)(fG)(mA)(fC)(mU)(fA)(mC)(fC)(mA)#(fA)#(mA)#(fG)#(mG)#(fC)#(mU)#(fC) | (fU)#(mU)#(fU)(mG)(fG)(mU)(fA)(mG)(fU)(mC)(fU)(mG)(fC)#(mG)#(fA)-TegChol | | | rs362307 | S5 |
| SNP 3-11 | 3 | 11 | P(mU)#(fC)#(mG)(fC)(mA)(fG)(mA)(fC)(mU)(fU)(mU)(fC)(mA)#(fA)#(mA)#(fG)#(mG)#(fC)#(mU)#(fC) | (fU)#(mU)#(fU)(mG)(fA)(mA)(fA)(mG)(fU)(mC)(fU)(mG)(fC)#(mG)#(fA)-TegChol | | | rs362307 | S5 |
| SNP 3-12 | 3 | 12 | P(mU)#(fC)#(mG)(fC)(mA)(fG)(mA)(fC)(mU)(fU)(mC)(fU)(mA)#(fA)#(mA)#(fG)#(mG)#(fC)#(mU)#(fC) | (fU)#(mU)#(fU)(mA)(fG)(mA)(fA)(mG)(fU)(mC)(fU)(mG)(fC)#(mG)#(fA)-TegChol | | | rs362307 | S5 |
| SNP 3-13 | 3 | 13 | P(mU)#(fC)#(mG)(fC)(mA)(fG)(mA)(fC)(mU)(fU)(mC)(fC)(mC)#(fA)#(mA)#(fG)#(mG)#(fC)#(mU)#(fC) | (fU)#(mU)#(fG)(mG)(fG)(mA)(fA)(mG)(fU)(mC)(fU)(mG)(fC)#(mG)#(fA)-TegChol | | | rs362307 | S5 |
| SNP 3-13G | 3 | 13 | P(mU)#(fC)#(mG)(fC)(mA)(fG)(mA)(fC)(mU)(fU)(mC)(fC)(mG)#(fA)#(mA)#(fG)#(mG)#(fC)#(mU)#(fC) | (fU)#(mU)#(fC)(mG)(fG)(mA)(fA)(mG)(fU)(mC)(fU)(mG)(fC)#(mG)#(fA)-TegChol | | | rs362307 | S5 |
| SNP 3-14C | 3 | 14 | P(mU)#(fC)#(mG)(fC)(mA)(fG)(mA)(fC)(mU)(fU)(mC)(fC)(mA)#(fC)#(mA)#(fG)#(mG)#(fC)#(mU)#(fC) | (fU)#(mG)#(fU)(mG)(fG)(mA)(fA)(mG)(fU)(mC)(fU)(mG)(fC)#(mG)#(fA)-TegChol | | | rs362307 | S5 |
| SNP 3-14G | 3 | 14 | P(mU)#(fC)#(mG)(fC)(mA)(fG)(mA)(fC)(mU)(fU)(mC)(fC)(mA)#(fG)#(mA)#(fG)#(mG)#(fC)#(mU)#(fC) | (fU)#(mC)#(fU)(mG)(fG)(mA)(fA)(mG)(fU)(mC)(fU)(mG)(fC)#(mG)#(fA)-TegChol | | | rs362307 | S5 |
| SNP 3-15C | 3 | 15 | P(mU)#(fC)#(mG)(fC)(mA)(fG)(mA)(fC)(mU)(fU)(mC)(fC)(mA)#(fA)#(mC)#(fG)#(mG)#(fC)#(mU)#(fC) | (fG)#(mU)#(fU)(mG)(fG)(mA)(fA)(mG)(fU)(mC)(fU)(mG)(fC)#(mG)#(fA)-TegChol | | | rs362307 | S5 |
| SNP 3-15G | 3 | 15 | P(mU)#(fC)#(mG)(fC)(mA)(fG)(mA)(fC)(mU)(fU)(mC)(fC)(mA)#(fA)#(mG)#(fG)#(mG)#(fC)#(mU)#(fC) | (fC)#(mU)#(fU)(mG)(fG)(mA)(fA)(mG)(fU)(mC)(fU)(mG)(fC)#(mG)#(fA)-TegChol | | | rs362307 | S5 |
| SNP6-11(A) Dio | 6 | 11 | V(mU)#(fU)#(mC)(fU)(mG)(fU)(mA)(fG)(mC)(fA)(mU)(fC)(mA)#(fG)#(mC)#(fU)#(mU)#(fC)#(mU)#(fC) | (fG)#(mC)#(fU)(mG)(fA)(mU)(fG)(mC)(fU)(mA)(fC)(mA)(fG)#(mA)#(fA)-Dio | | | rs362273 | 5,S6,S7,S8 |

| notation | meaning |
|---|---|
| P | phosphate group |
| V | vinylphosphonate |
| m | 2'Ome |
| f | 2'F |
| # | phosphorothioate |
| TegChol | teg linker linked to cholesterol |
| Dio | divalent siRNAs for in vivo use have a teg linker attached to an additional sense strand, no cholesterol. |

| siRNA ID | SNP position | additional mismatch position | ntisense strand sequen | sense strand sequence | IC 50 target [nM] | IC50 non-target [nM] | Target SNP site |
|---|---|---|---|---|---|---|---|
| SNP2-0 | 2 | 0 | UUAGCAGCAGCUUCUCGUGG | AGAAGCUGCUGCUAA | | | rs362273 |
| SNP2-3G | 2 | 3 | UUGGCAGCAGCUUCUCGUGG | AGAAGCUGCUGCUAA | | | rs362273 |
| SNP2-3U | 2 | 3 | UUUGCAGCAGCUUCUCGUGG | AGAAGCUGCUGCUAA | | | rs362273 |
| SNP2-3C | 2 | 3 | UUCGCAGCAGCUUCUCGUGG | AGAAGCUGCUGCUAA | | | rs362273 |
| SNP2-4 | 2 | 4 | UUAUCAGCAGCUUCUCGUGG | AGAAGCUGCUGCUAA | | | rs362273 |
| SNP2-5 | 2 | 5 | UUAGUAGCAGCUUCUCGUGG | AGAAGCUGCUGCUAA | | | rs362273 |
| SNP2-6 | 2 | 6 | UUAGCUGCAGCUUCUCGUGG | AGAAGCUGCUGCUAA | | | rs362273 |
| SNP2-7 | 2 | 7 | UUAGCAUCAGCUUCUCGUGG | AGAAGCUGCUGCUAA | | | rs362273 |
| SNP2-8 | 2 | 8 | UUAGCAGUAGCUUCUCGUGG | AGAAGCUGCUGCUAA | | | rs362273 |
| SNP2-9 | 2 | 9 | UUAGCAGCUGCUUCUCGUGG | AGAAGCUGCUGCUAA | | | rs362273 |
| SNP2-10 | 2 | 10 | UUAGCAGCAUCUUCUCGUGG | AGAAGCUGCUGCUAA | | | rs362273 |
| SNP2-11 | 2 | 11 | UUAGCAGCAGUUUCUCGUGG | AGAAGCUGCUGCUAA | | | rs362273 |
| SNP2-12 | 2 | 12 | UUAGCAGCAGCAUCUCGUGG | AGAAGCUGCUGCUAA | | | rs362273 |
| SNP2-13 | 2 | 13 | UUAGCAGCAGCUACUCGUGG | AGAAGCUGCUGCUAA | | | rs362273 |
| SNP2-14 | 2 | 14 | UUAGCAGCAGCUUUUCGUGG | AGAAGCUGCUGCUAA | | | rs362273 |
| SNP2-15 | 2 | 15 | UUAGCAGCAGCUUCACGUGG | AGAAGCUGCUGCUAA | | | rs362273 |
| SNP2-16 | 2 | 16 | UUAGCAGCAGCUUCUUGUGG | AGAAGCUGCUGCUAA | | | rs362273 |
| SNP3-0 | 3 | 0 | UGUAGCAGCAGCUUCUCGUG | GAAGCUGCUGCUACA | | | rs362273 |
| SNP4-0 | 4 | 0 | UUGUAGCAGCAGCUUCUCGU | AAGCUGCUGCUACAA | 38 | 803 | rs362273 |
| SNP4-2 | 4 | 2 | UAGUAGCAGCAGCUUCUCGU | AAGCUGCUGCUACAA | | | rs362273 |
| SNP4-3U | 4 | 3 | UUUUAGCAGCAGCUUCUCGU | AAGCUGCUGCUACAA | | | rs362273 |
| SNP4-3A | 4 | 3 | UUAUAGCAGCAGCUUCUCGU | AAGCUGCUGCUACAA | | | rs362273 |
| SNP4-5C | 4 | 5 | UUGUCGCAGCAGCUUCUCGU | AAGCUGCUGCUACAA | | | rs362273 |
| SNP4-5G | 4 | 5 | UUGUGGCAGCAGCUUCUCGU | AAGCUGCUGCUACAA | | | rs362273 |
| SNP4-6 | 4 | 6 | UUGUAUCAGCAGCUUCUCGU | AAGCUGCUGCUACAA | | | rs362273 |
| SNP4-7 | 4 | 7 | UUGUAGUAGCAGCUUCUCGU | AAGCUGCUGCUACAA | 17 | >5000 | rs362273 |
| SNP4-8 | 4 | 8 | UUGUAGCUGCAGCUUCUCGU | AAGCUGCUGCUACAA | 24 | 220 | rs362273 |
| SNP4-9 | 4 | 9 | UUGUAGCAUCAGCUUCUCGU | AAGCUGCUGCUACAA | | | rs362273 |
| SNP4-10 | 4 | 10 | UUGUAGCAGUAGCUUCUCGU | AAGCUGCUGCUACAA | | | rs362273 |
| SNP4-11 | 4 | 11 | UUGUAGCAGCUGCUUCUCGU | AAGCUGCUGCUACAA | | | rs362273 |
| SNP4-12 | 4 | 12 | UUGUAGCAGCAUCUUCUCGU | AAGCUGCUGCUACAA | | | rs362273 |
| SNP4-13 | 4 | 13 | UUGUAGCAGCAGUUUCUCGU | AAGCUGCUGCUACAA | | | rs362273 |

| | | | | | | | |
|---|---|---|---|---|---|---|---|
| SNP4-14 | 4 | 14 | UUGUAGCAGCAGCAUCUCGU | AAGCUGCUGCUACAA | | | rs362273 |
| SNP4-15 | 4 | 15 | UUGUAGCAGCAGCUACUCGU | AAGCUGCUGCUACAA | | | rs362273 |
| SNP4-16 | 4 | 16 | UUGUAGCAGCAGCUUUUCGU | AAGCUGCUGCUACAA | | | rs362273 |
| SNP5-0 | 5 | 0 | UCUGUAGCAGCAGCUUCUCG | AGCUGCUGCUACAGA | | | rs362273 |
| SNP6-0 | 6 | 0 | UUCUGUAGCAGCAGCUUCUC | GCUGCUGCUACAGAA | 24 | 174 | rs362273 |
| SNP6-2 | 6 | 2 | UACUGUAGCAGCAGCUUCUC | GCUGCUGCUACAGAA | | | rs362273 |
| SNP6-3 | 6 | 3 | UUUUGUAGCAGCAGCUUCUC | GCUGCUGCUACAGAA | | | rs362273 |
| SNP6-4 | 6 | 4 | UUCAGUAGCAGCAGCUUCUC | GCUGCUGCUACAGAA | | | rs362273 |
| SNP6-5U | 6 | 5 | UUCUUUAGCAGCAGCUUCUC | GCUGCUGCUACAGAA | | | rs362273 |
| SNP6-5A | 6 | 5 | UUCUAUAGCAGCAGCUUCUC | GCUGCUGCUACAGAA | | | rs362273 |
| SNP6-7C | 6 | 7 | UUCUGUCGCAGCAGCUUCUC | GCUGCUGCUACAGAA | | | rs362273 |
| SNP6-7G | 6 | 7 | UUCUGUGGCAGCAGCUUCUC | GCUGCUGCUACAGAA | | | rs362273 |
| SNP6-8 | 6 | 8 | UUCUGUAUCAGCAGCUUCUC | GCUGCUGCUACAGAA | | | rs362273 |
| SNP6-9 | 6 | 9 | UUCUGUAGUAGCAGCUUCUC | GCUGCUGCUACAGAA | | | rs362273 |
| SNP6-10 | 6 | 10 | UUCUGUAGCUGCAGCUUCUC | GCUGCUGCUACAGAA | | | rs362273 |
| SNP6-11 (G) | 6 | 11 | UUCUGCAGCAUCAGCUUCUC | GCUGCUGCUGCAGAA | | | rs362273 |
| SNP6-11 (A) | 6 | 11 | UUCUGUAGCAUCAGCUUCUC | GCUGCUGCUACAGAA | 28 | >4500 | rs362273 |
| SNP6-12 | 6 | 12 | UUCUGUAGCAGUAGCUUCUC | GCUGCUGCUACAGAA | 24 | >1000 | rs362273 |
| SNP6-13 | 6 | 13 | UUCUGUAGCAGCUGCUUCUC | GCUGCUGCUACAGAA | | | rs362273 |
| SNP6-14 | 6 | 14 | UUCUGUAGCAGCAUCUUCUC | GCUGCUGCUACAGAA | 30 | >2000 | rs362273 |
| SNP6-15 | 6 | 15 | UUCUGUAGCAGCAGUUUCUC | GCUGCUGCUACAGAA | | | rs362273 |
| SNP6-16 | 6 | 16 | UUCUGUAGCAGCAGCAUCUC | GCUGCUGCUACAGAA | 18 | 333 | rs362273 |
| SNP7-0 | 7 | 0 | UAUCUGUAGCAGCAGCUUCU | CUGCUGCUACAGAUA | | | rs362273 |
| SNP8-0 | 8 | 0 | UGAUCUGUAGCAGCAGCUUC | UGCUGCUACAGAUCA | | | rs362273 |
| SNP9-0 | 9 | 0 | UUGAUCUGUAGCAGCAGCUU | GCUGCUACAGAUCAA | | | rs362273 |
| SNP10-0 | 10 | 0 | UUUGAUCUGUAGCAGCAGCU | CUGCUACAGAUCAAA | | | rs362273 |
| SNP11-0 | 11 | 0 | UGUUGAUCUGUAGCAGCAGC | UGCUACAGAUCAACA | | | rs362273 |
| SNP12-0 | 12 | 0 | UGGUUGAUCUGUAGCAGCAG | GCUACAGAUCAACCA | | | rs362273 |
| SNP13-0 | 13 | 0 | UGGGUUGAUCUGUAGCAGCA | CUACAGAUCAACCCA | | | rs362273 |
| SNP2 | 2 | 0 | UGCAGACUUCCAAAGGCUCC | CUUUGGAAGUCUGCA | | | rs362307 |
| SNP3 | 3 | 0 | UCGCAGACUUCCAAAGGCUC | UUUGGAAGUCUGCGA | 35 | 438 | rs362307 |
| SNP4 | 4 | 0 | UGCGCAGACUUCCAAAGGCU | UUGGAAGUCUGCGCA | | | rs362307 |
| SNP5 | 5 | 0 | UGGCGCAGACUUCCAAAGGC | UGGAAGUCUGCGCCA | | | rs362307 |
| SNP6 | 6 | 0 | UGGGCGCAGACUUCCAAAGG | GGAAGUCUGCGCCCA | | | rs362307 |
| SNP7 | 7 | 0 | UAGGGCGCAGACUUCCAAAG | GAAGUCUGCGCCCUA | | | rs362307 |

| | | | | | | | | |
|---|---|---|---|---|---|---|---|---|
| SNP8 | 8 | 0 | | UAAGGGCGCAGACUUCCAAA | AAGUCUGCGCCCUUA | | | rs362307 |
| SNP9 | 9 | 0 | | UCAAGGGCGCAGACUUCCAA | AGUCUGCGCCCUUGA | | | rs362307 |
| SNP10 | 10 | 0 | | UACAAGGGCGCAGACUUCCA | GUCUGCGCCCUUGUA | | | rs362307 |
| SNP11 | 11 | 0 | | UCACAAGGGCGCAGACUUCC | UCUGCGCCCUUGUGA | | | rs362307 |
| SNP12 | 12 | 0 | | UGCACAAGGGCGCAGACUUC | CUGCGCCCUUGUGCA | | | rs362307 |
| SNP13 | 13 | 0 | | UGGCACAAGGGCGCAGACUU | UGCGCCCUUGUGCCA | | | rs362307 |
| SNP 3-2 | 3 | 2 | | UUGCAGACUUCCAAAGGCUC | UUUGGAAGUCUGCAA | | | rs362307 |
| SNP 3-4 | 3 | 4 | | UCGUAGACUUCCAAAGGCUC | UUUGGAAGUCUACGA | | | rs362307 |
| SNP 3-5C | 3 | 5 | | UCGCCGACUUCCAAAGGCUC | UUUGGAAGUCGGCGA | >1500 | 92 | rs362307 |
| SNP 3-5G | 3 | 5 | | UCGCGGACUUCCAAAGGCUC | UUUGGAAGUCCGCGA | | | rs362307 |
| SNP 3-6 | 3 | 6 | | UCGCAUACUUCCAAAGGCUC | UUUGGAAGUAUGCGA | | | rs362307 |
| SNP 3-7C | 3 | 7 | | UCGCAGCCUUCCAAAGGCUC | UUUGGAAGGCUGCGA | 400 | 1500 | rs362307 |
| SNP 3-7G | 3 | 7 | | UCGCAGGCUUCCAAAGGCUC | UUUGGAAGCCUGCGA | 20 | 83 | rs362307 |
| SNP 3-8 | 3 | 8 | | UCGCAGAUUUCCAAAGGCUC | UUUGGAAAUCUGCGA | 20 | 750 | rs362307 |
| SNP 3-9 | 3 | 9 | | UCGCAGACAUCCAAAGGCUC | UUUGGAUGUCUGCGA | | | rs362307 |
| SNP 3-10 | 3 | 10 | | UCGCAGACUACCAAAGGCUC | UUUGGUAGUCUGCGA | | | rs362307 |
| SNP 3-11 | 3 | 11 | | UCGCAGACUUUCAAAGGCUC | UUUGAAAGUCUGCGA | | | rs362307 |
| SNP 3-12 | 3 | 12 | | UCGCAGACUUCUAAAGGCUC | UUUAGAAGUCUGCGA | | | rs362307 |
| SNP 3-13 | 3 | 13 | | UCGCAGACUUCCCAAGGCUC | UUGGGAAGUCUGCGA | | | rs362307 |
| SNP 3-13G | 3 | 13 | | UCGCAGACUUCCGAAGGCUC | UUCGGAAGUCUGCGA | | | rs362307 |
| SNP 3-14C | 3 | 14 | | UCGCAGACUUCCACAGGCUC | UGUGGAAGUCUGCGA | | | rs362307 |
| SNP 3-14G | 3 | 14 | | UCGCAGACUUCCAGAGGCUC | UCUGGAAGUCUGCGA | | | rs362307 |
| SNP 3-15C | 3 | 15 | | UCGCAGACUUCCAACGGCUC | GUUGGAAGUCUGCGA | | | rs362307 |
| SNP 3-15G | 3 | 15 | | UCGCAGACUUCCAAGGGCUC | CUUGGAAGUCUGCGA | | | rs362307 |
